# The volatile cedrene from plant beneficial *Trichoderma guizhouense* modulates *Arabidopsis* root development through auxin transport and signaling

**DOI:** 10.1101/2021.04.09.439204

**Authors:** Yucong Li, Jiahui Shao, Yansong Fu, Yu Chen, Hongzhe Wang, Zhihui Xu, Haichao Feng, Weibing Xun, Yunpeng Liu, Nan Zhang, Qirong Shen, Wei Xuan, Ruifu Zhang

**Author notes:** **Corresponding authors:** Ruifu Zhang.; Wei Xuan. These authors contributed equally to this work (Yucong Li, Jiahui Shao). **Footnotes:** This work was funded by the National Natural Science Foundation of China (31972512 and 32072665), the Agricultural Science and Technology Innovation Program of CAAS (No. CAAS-ZDRW202009) and Key R&D Program of Shandong Province (2019JZZY020614). **Author contributions:** R.Z. planned and designed the research. Y.C.L. performed the experiments and wrote the manuscript. R.Z. and W.X. polished the manuscript. J.H.S. and Y.S.F. and helped with root phenotype analysis and imaging. Y.C. and H.Z.W. analyzed the data. All authors discussed the results. R.Z. and W.X. agree to serve as the authors responsible for contact and ensure communication.

## Abstract

Rhizosphere microorganisms interact with plant roots by producing chemical signals to regulate root development. However, the involved distinct bioactive compounds and the signal transduction pathways are remaining to be identified. Here, we show that sesquiterpenes (SQTs) are the main volatile compounds produced by plant beneficial *Trichoderma guizhouense* NJAU 4742, inhibition of SQTs synthesis in this strain indicated their involvement in plant-fungus cross-kingdom signaling. SQTs component analysis further identified the cedrene, a high abundant SQT in strain NJAU 4742, could stimulate plant growth and root development. Genetic analysis and auxin transport inhibition showed that auxin receptor TIR1, AFB2, auxin-responsive protein IAA14, and transcription factor ARF7, ARF19 affect the response of lateral roots to cedrene. Moreover, auxin influx carrier AUX1, efflux carrier PIN2 were also indispensable for cedrene-induced lateral root formation. Confocal imaging showed that cedrene affected the expression of *pPIN2:PIN2:GFP* and *pPIN3:PIN3:GFP*, which may be related to the effect of cedrene on root morphology. These results suggest that a novel SQT molecule from plant beneficial *T. guizhouense* can regulate plant root development through auxin transport and signaling.

**One-sentence Summary:** Cedrene, a high- abundance sesquiterpenes produced by plant beneficial *Trichoderma guizhouense* NJAU 4742, stimulates *Arabidopsis* lateral root formation and primary root elongation by relying on auxin signaling pathway and auxin transporter PIN2 and AUX1.

## INTRODUCTIONS

The periodic formation of lateral roots (LR) is a post-embryonic process that is regulated by both endogenous and environmental cues (Xie et al., 2019; Li et al., 2021). Auxin plays the central role in all stages of LR formation, which consists of LR priming, LR initiation and patterning and LR emergence (Lavenus et al., 2013; Santos Teixeira and Ten Tusscher, 2019). Following the act of lateral root cap (LRC) derived auxin on the oscillation zone (OZ), an oscillatory signal is generated to control xylem pole pericycle cells (XPP) ‘priming’ and pre-branch sites forming (Xuan et al., 2020). Auxin regulates LR initiation by controlling LR founder cell divisions and LR founder cell polarity and/or identity acquisition (Dubrovsky et al., 2008). In the LR emergence stage, auxin induces the expression of cell wall-remodeling enzymes to promote cell separation, changes the cell turgor pressure in the outer tissue layers and in the lateral root primordia (LRP), these progresses will help LRPs to go through three overlaying cell layers (endodermis, cortex, and epidermis) to emerge as the LR (Neuteboom et al., 1999; Laskowski et al., 2006; Swarup et al., 2008; Peret et al., 2012; Lee et al., 2013).

Environmental factors, which include water, salt, drought, light, nitrate and phosphate, affect LR formation by interfering with auxin synthesis, conjugation, and degradation, as well as auxin transport or response (Santos Teixeira and Ten Tusscher, 2019).The rhizosphere, defined as the narrow zone influenced by plant roots and characterized by their intense association with microbial activity (Mendes et al., 2013; van Dam and Bouwmeester, 2016), is relatively rich in nutrients, because about 20-40% of the photosynthetic products can be lost from the root in the form of root exudates, including ions, oxygen, water, enzymes, mucus and primary and secondary metabolites (Bais et al., 2006; Philippot et al., 2013; Venturi and Keel, 2016). Consequently, a large number and variety of microorganisms, including bacteria and fungi, were inhabited in the rhizosphere. The rhizosphere microorganisms affect root system architecture (RSA) by the production of phytohormones and secondary metabolites, such as auxin, cytokinin and 2,4-diacetylphloroglucinol (DAPG), which interfere with auxin-dependent signaling pathways in plants (Vacheron et al., 2013). Microbes also produce many volatile compounds (VCs) with the ability to reprogram RSA (Zhang et al., 2007; Kanchiswamy et al., 2015; Werner et al., 2016; Tyc et al., 2017; Fincheira and Quiroz, 2018), which can evaporate and diffuse easily far from their original point and migrate in soil and aerial environments for their low molecular masses and low polarity, and as well as a high vapor pressure (Schulz and Dickschat, 2007; Schmidt et al., 2015).VC component of indole emitted by soil-borne bacteria affected LR development in *Arabidopsis* through auxin signaling, depend on polar auxin transport system (Bailly et al., 2014). (–)-Thujopsene, a kind of sesquiterpenes (SQTs), produced by *Laccaria bicolor*, stimulated LR formation in *Arabidopsis* by inducing superoxide anion radicals in roots, independent on auxin signaling pathways (Ditengou et al., 2015). 6-pentyl-2H-pyran-2-one (6-PP) detected in *Trichoderma atroviride* regulated LR development through auxin signaling and transport in the root system (Garnica-Vergara et al., 2016). The VCs produced *by T. viride* with the main ingredients of isobutyl alcohol, isopentyl alcohol, and 3-methylbutanal, increased biomass in the shoot and the root system (Hung et al., 2013). So far, considerable progress has been made in elucidating the mode of action of VCs; however, it is still poorly understood, especially in the identification of distinct bioactive compounds.

*Trichoderma* species are able to colonize the root surface to promote the root development and plant growth, which also represent excellent biocontrol agents in agriculture because of their strong ability to fight with plant pathogens (Druzhinina et al., 2011). *Trichoderma* species have broad VC profiles (Hung et al., 2013; Muller et al., 2013), the produced SQTs were supposed to be good candidates for underground microbe-plant signaling as SQTs were the representative diffusing compounds in complex environment (Hiltpold and Turlings, 2008). In this study, we identified the cedrene from *Trichoderma guizhouense* NJAU 4742, the plant beneficial fungus with efficient promotion for plant growth and root development, as well as the soil-borne pathogen suppression (Zhang et al., 2016; Meng et al., 2019; Zhang et al., 2019). The cedrene can enhance shoot and root biomass and stimulate root development in a dose-dependent manner. Cedrene induced auxin response in primary root tips and early stages LRP (I-III), but not the late stages LRP (IV-VII). Furthermore, cedrene differentially modulated expression of auxin transporters PIN1, PIN2, PIN3 and PIN7. A genetic screen for cedrene resistance established that this compound required auxin receptors TIR1 and AFB2, and downstream auxin-responsive protein IAA14, and transcription factors ARF7 and ARF19 to stimulate lateral root development. Moreover, auxin influx carrier AUX1 and efflux carrier PIN2 were both indispensable for promoting lateral root formation.

## RESULTS

### VCs released by *T. guizhouense* NJAU 4742 promote *Arabidopsis* growth and root development

*Arabidopsis* plants were grown in the presence of fungal VCs in bi-compartmented Petri dishes (Fig. 1a). After 8 d of co-cultivation, despite the absence of direct contact with *T. guizhouense* NJAU 4742, strong stimulation of growth and root development was observed in *Arabidopsis*, which should have been caused by VCs of NJAU 4742. *T. guizhouense* VCs slightly promoted primary root elongation, but significantly increased the lateral root number and lateral root density by 37% and 39%, respectively (Fig.1a-c). In addition, *T. guizhouense* VCs can also increase the root and shoot biomass by 101% and 56% (Fig.1e, f), respectively, and increase the total biomass production of plants by 71% (Fig.1g).

**Figure 1.**
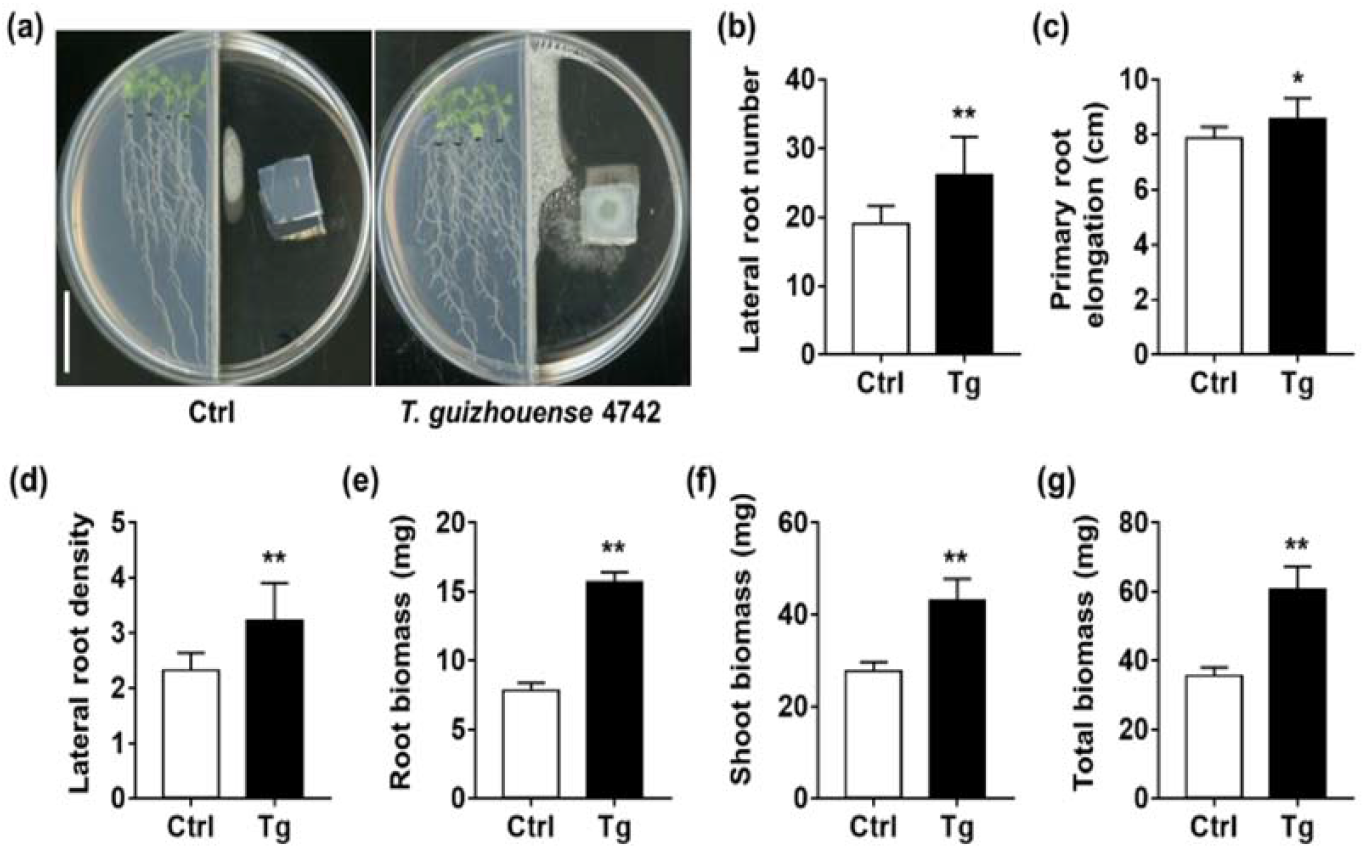
Effect of *Trichoderma guizhouense* NJAU 4742 volatile compounds on *Arabidopsis* biomass production and root development. (a) Three-day-old *Arabidopsis thaliana* (Col-0) seedlings were germinated and grown for 8 d in the presence of *T. guizhouense* NJAU 4742 (Tg) in a bi-compartmented Petri dish avoiding direct contact and solute exchange between the plant and the fungus. Scale bar, 2 cm. (b and c) Quantification of primary root elongation (b) and number of emerged lateral roots (c). n = 12. **, P < 0.01; *, P < 0.05 (Student’s *t*-test). (d) Lateral root density (number of emerged lateral roots cm^-1^). n = 12. **, P < 0.01; *, P < 0.05 (Student’s *t*-test). (e-g) Quantification of root biomass (e), shoot biomass (f) and total biomass (g) production per *Arabidopsis* seedling. n = 12. **, P < 0.01; *, P < 0.05 (Student’s *t*-test). Error bars indicate ± SD of the mean.

### Cedrene mimics fungal VCs effects in promoting LR formation

To identify the VCs component responsible for LR induction, VCs of *T. guizhouense* NJAU 4742 in the headspace was measured by SPME-GC-MS. Table 1 shows that sesquiterpenes (SQTs) are the major compounds within the VC profile (75.93%) from *T. guizhouense* NJAU 4742. Previous studies have shown that SQTs from ectomycorrhizal fungi *Laccaria bicolor* can reprogramme root architecture (Ditengou et al., 2015). To test whether SQTs in *T. guizhouense* VCs are related to the induction of lateral root formation, we inhibited SQT biosynthesis using inhibitor lovastatin (Rodriguez-Concepcion, 2006; Ditengou et al., 2015) in *T. guizhouense* NJAU 4742 and investigated the effect on LR stimulation in *Arabidopsis*. The result showed that the effect of *T. guizhouense* VCs on promoting root development was abolished when fungal SQTs were suppressed by lovastatin (Fig. 2a-c). To identify distinct bioactive compounds, we tested the effect of a pure product (cedrene) that can be obtained and contained in a large amount in *T. guizhouense* VCs profile on the plant growth-promoting activity. The *Arabidopsis* seedlings were treated with DMSO (as control) or with 10-2000 μM cedrene dissolved in DMSO (Fig. 3a). After 8 d of growth in medium supplied with 20-100 μM cedrene, a significant increase in shoot, root and total plant biomass was observed (Fig. 3b-d). By contrast, the higher concentration (250-1000 μM) or lower concentration (10 μM) of cedrene had no effect on the growth, and the highest concentration (2000 μM) even inhibited the growth of *Arabidopsis* (Fig. 3b-d). It is noteworthy that appropriate concentration (20-500 μM) and lower concentration (20-100 μM) of cedrene could induce the increase of lateral root number and primary root elongation, respectively (Fig. 3e, f), but higher concentration cedrene would have no or even inhibitory effect on that (Fig. 3e, f). Moreover, the cedrene treatments increased lateral root density in a dose-dependent manner (Fig. 3g).

**Table 1.** Volatile compounds produced by *Trichoderma guizhouense* NJAU 4742 after 6 d of growth in 1/2 MS salts agar medium, analyzed by solid-phase microextraction (SPME)-GC-MS

|  | Metabolite name | Normalized amount<br>of volatile<br>compound (%) |
| --- | --- | --- |
| Sesquiterpene<br>s | 1,4-Cadinadiene | 25.50 ± 4.08 |
|  | Cedrene | 17.11 ± 4.29 |
|  | Dauca-4(11),8-diene | 8.66 ± 2.69 |
|  | Cis-calamenene | 9.31 ± 4.54 |
|  | (+)-Cuparene | 4.28 ± 1.76 |
|  | Copaene | 3.57 ± 1.54 |
|  | (+)-Acoradiene | 2.39 ± 0.57 |
|  | g-Muurolene | 1.89 ± 0.35 |
|  | 4,9-Cadinadiene | 0.93 ± 0.11 |
|  | β-Cubebene | 0.74 ± 0.08 |
|  | Cycloisolongifolene | 0.70 ± 0.34 |
|  | Aromandendrene | 0.21 ± 0.09 |
|  | β-Curcumene | 0.19 ± 0.09 |
| other VCs | Benzene, 1,4-diethyl- | 17.24 ± 4.29 |
|  | 3,3-Dimethylphthalide | 1.59 ± 0.86 |
|  | D-alanine, n-(4-butylbenzoyl)-, isohexyl ester | 1.05 ± 0.16 |
|  | 4-hepten-3-one, 5-ethyl-4-methyl- | 1.31 ± 0.84 |
|  | 6,6-dimethyl-2-vinylidenebicyclo[3.1.1]heptane | 0.94 ± 0.71 |
|  | Spiro[4.5]dec-6-en-8-one,<br>1,7-dimethyl-4-(1-methylethyl)- | 0.52 ± 0.26 |
|  | Benzene, 1-(1,5-dimethylhexyl)-4-methyl- | 0.46 ± 0.14 |
|  | 1-bromo-3,7-dimethyl-2,6-octadiene | 0.32 ± 0.13 |
|  | Spiro[4.5]dec-8-en-7-ol,<br>4,8-dimethyl-1-(1-methylethyl)- | 0.26 ± 0.11 |

Normalized amount of volatile compound = (peak area of volatile compound) / (total peak area of all volatile compounds). Compounds were qualitative analyzed on the basis of NIST library (https://webbook.nist.gov/chemistry/). Values represent means of three independent determinations.

**Figure 2.**
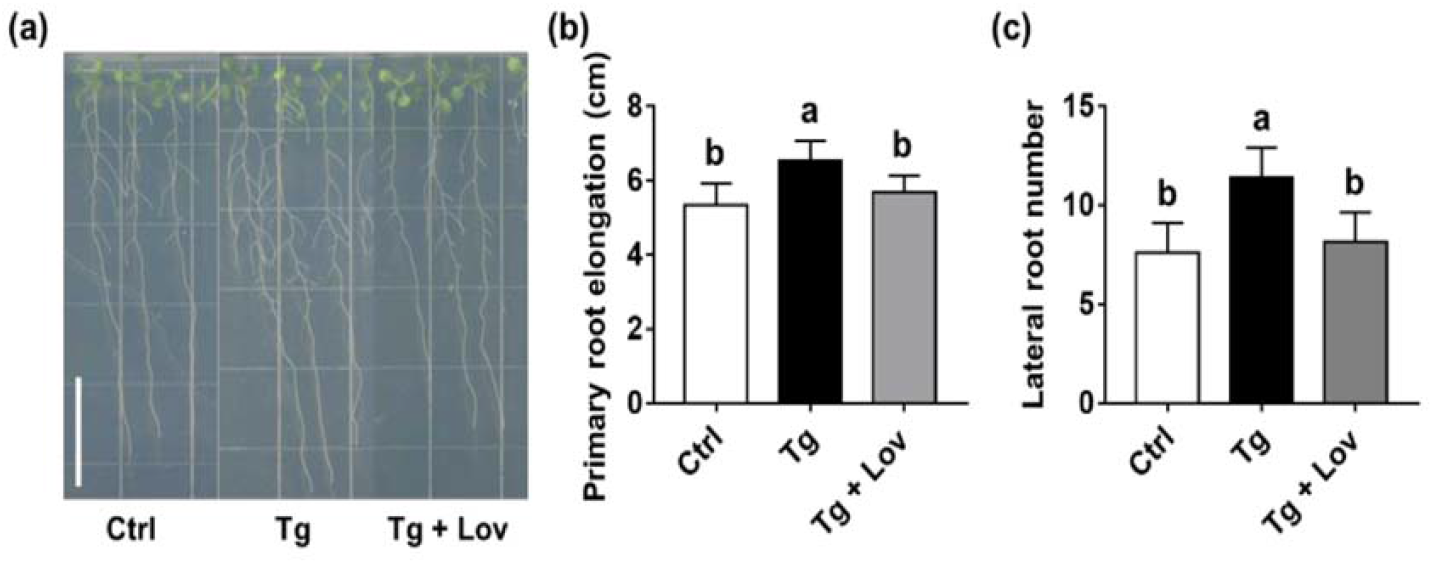
Sesquiterpenes in *Trichoderma guizhouense* NJAU 4742 volatile compounds are the main bioactive compounds for inducing *Arabidopsis* root development. (a) Three-day-old *Arabidopsis* seedlings were grown for 7 d in bi-compartmented Petri dishes (10cm × 10cm), which was inoculated with *T. guizhouense* in the presence (Tg+Lov) or absence (Tg) of 10 μM lovastatin (Lov) in the adjacent compartment. Scale bar, 2 cm. (b and c) Quantification of primary root elongation (b) and emerged lateral roots number of plants shown in (a). Different letters indicate significant differences of the mean values at P < 0.05 (One-way ANOVA, n = 12). Error bars indicate ± SD of the mean.

**Figure 3.**
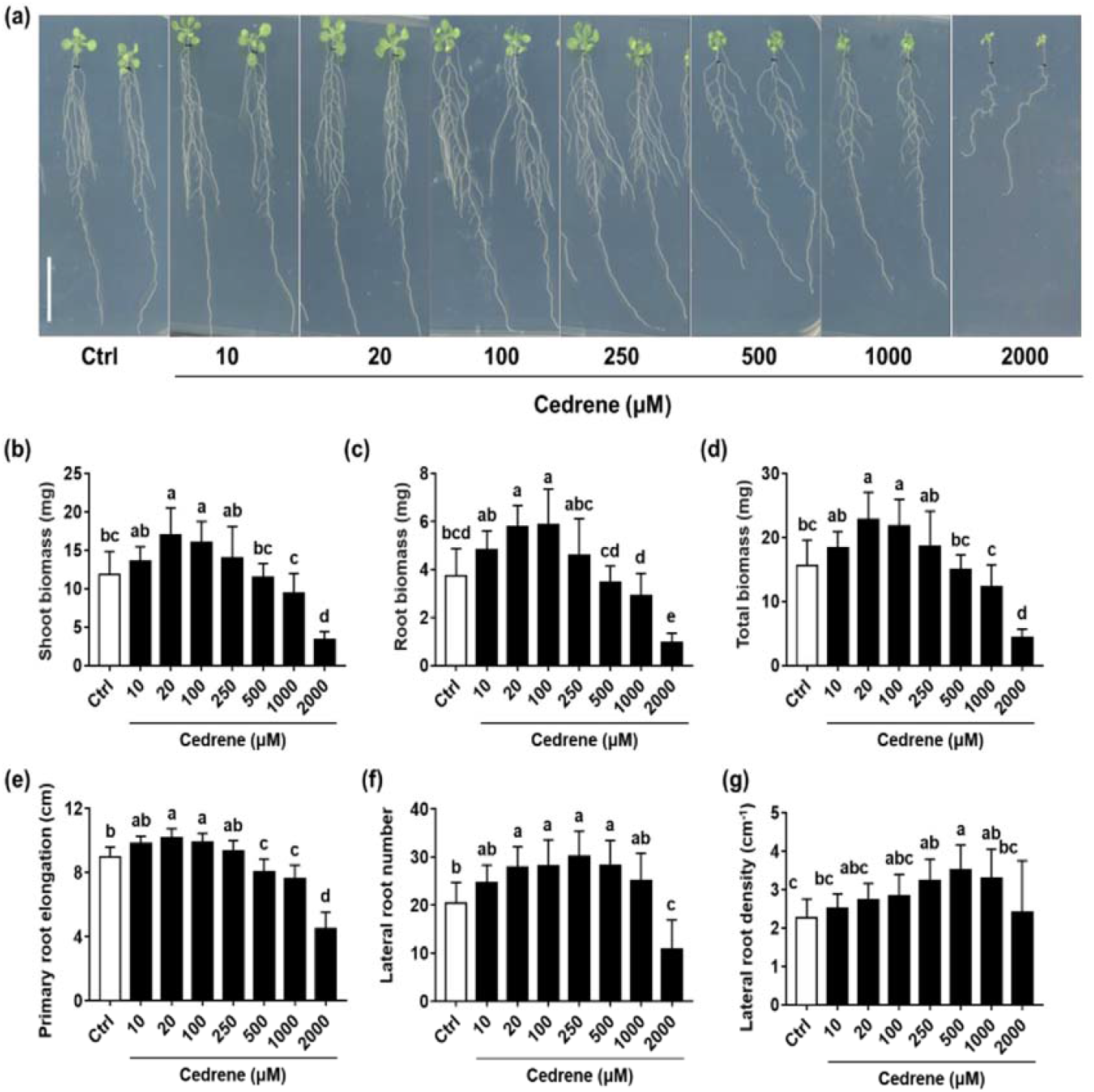
Effect of cedrene on *Arabidopsis* biomass production and root development. (a) Representative photographs of *Arabidopsis* seedlings grown for 8 d in 1/2 MS salts agar medium supplied with the solvent (DMSO) or increasing concentrations of cedrene. Scale bar, 2 cm. (b-d) Quantification of shoot biomass (b), root biomass (c), and total biomass (d) per seedling of plants shown in (a). Different letters indicate significant differences of the mean values at P < 0.05 (One-way ANOVA, n = 12). (e-g) Quantification of primary root elongation (e), emerged lateral roots number (f), and lateral root density (g, number of emerged lateral roots cm^-1^) of plants shown in (a). Different letters indicate significant differences of the mean values at P < 0.05 (One-way ANOVA, n = 12). Error bars indicate ± SD of the mean.

### Cedrene affects early LRP stages to induce lateral root formation

To further investigate the stage of LRP that is impacted by cedrene treatment, the developmental stage of LRP in the newly formed PR of control and cedrene-treated plants were quantified. Cedrene induced more LRP formation at very early stages (Stages I) compared with control treatment (Fig. 4c). However, we did not observe changes on later stages LRP (from Stage II to Stage VII) in cedrene-treated seedling roots (Fig. 4c). It is noteworthy that cedrene treatment did not affect the total LRP number (Fig. 4c). Next, we analyzed auxin distribution using the *pDR5:GUS* reporter and an enhanced DR5-directed GUS activity was detected in early stages (Stages I to III) LRP and meristematic protoxylem pole (Fig. 4a, b). Moreover, a strong and diffuse increase of *GUS* activity in primary root of auxin-inducible LRP specific marker line *pGATA23:nls-GUS* (De Rybel et al., 2010) was observed when treated with cedrene (Fig. 4d). Together, these results demonstrated that cedrene might stimulate early lateral roots initiation and following LRP development by affecting auxin distribution and its downstream signaling.

**Figure 4.**
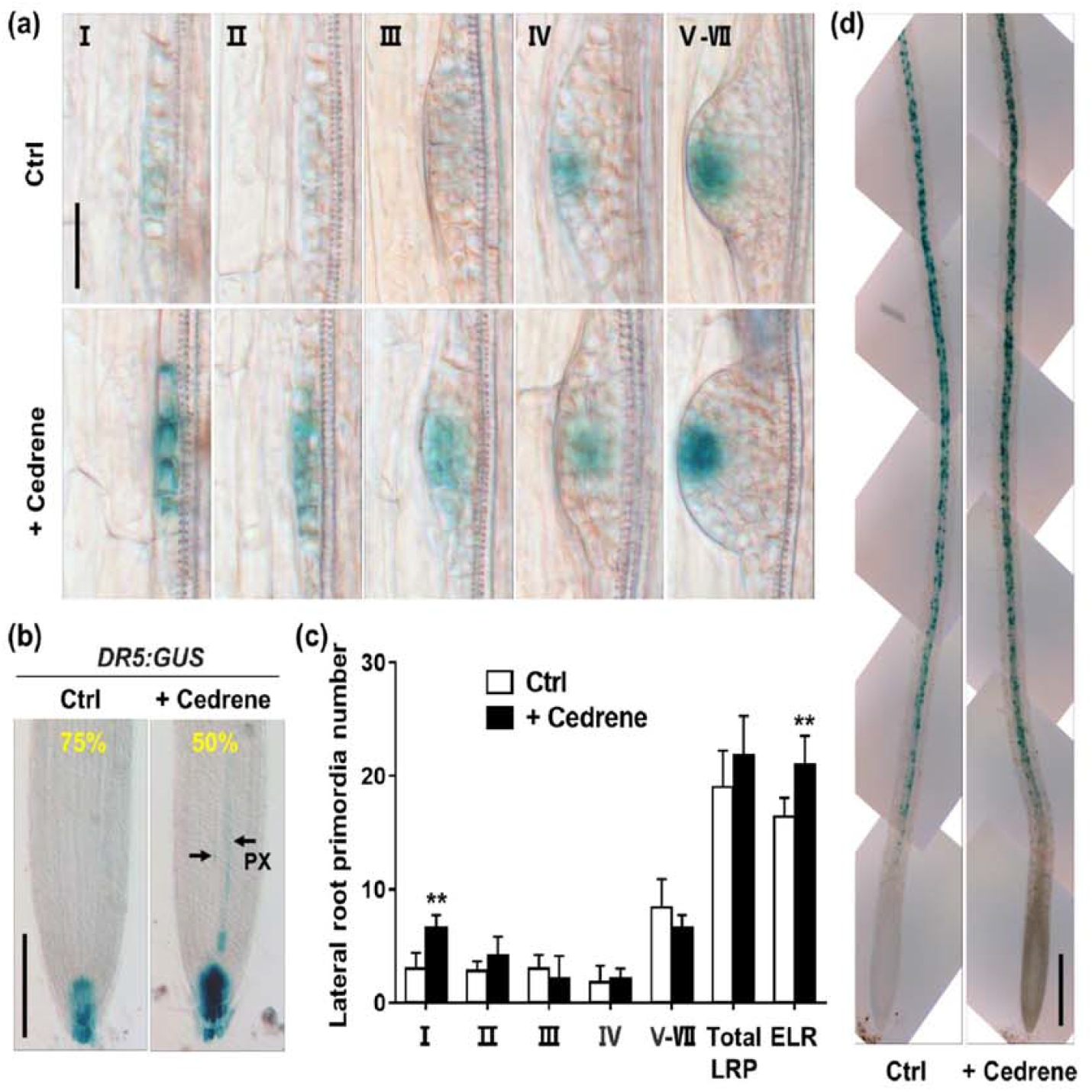
The effects of cedrene on lateral root primordium development in Arabidopsis. (a) Different stages of lateral root primordia expressing *pDR5:GUS* under solvent (DMSO, Ctrl) or 100 μM cedrene-treated conditions. Scale bar, 100 μm. (b) Expression pattern of *DR5:GUS* in the root tips of 3-day-old seedling treated with or without cedrene for six more days. Percentages indicate the proportion of seedlings showing the identical *DR5* expression pattern within a population (n = 12). PX, protoxylem pole. Scale bar, 100 μm. (c) Distribution of lateral root primordia (LRP) in seven developmental classes as defined by the *pCYCB1;1:GUS* activity after treatment with 100 μM cedrene for 7 more days. n = 8. **, P < 0.01; *, P < 0.05 (Student’s *t*-test). Total LRP, total number of lateral root primordia including all seven developmental stages. ELR, emerged lateral roots. (d) *pGATA23:nls-GUS* expression in primary root under control and cedrene-treated (100 μM) conditions. Scale bar, 200 μm.

### Cedrene does not alter *DR5* oscillation in roots

In order to investgate whether cedrene is involved in the regulation of ‘lateral root clock’, which generates an oscillatory signal that is translated into a developmental cue to specify a set of founder cells for LR formation (Xuan *et al*., 2020), we used *pDR5:LUC* (Fig. 5a), which fused *DR5* promoter to the *luciferase* (LUC) gene to mark early LR founder cell positions and allow visualization of its behavior *in vivo* (Moreno-Risueno et al., 2010; Van Norman et al., 2013; Laskowski and Ten Tusscher, 2017; Xuan et al., 2020). Expression of *pDR5:LUC* in the root tip showed oscillatory activity, and a static point of expression, which is the future site of LRP and LR and therefore is called as prebranch sites (Moreno-Risueno *et al*., 2010; Van Norman *et al*., 2013) was formed following each peak of the *DR5* oscillation.

**Figure 5.**
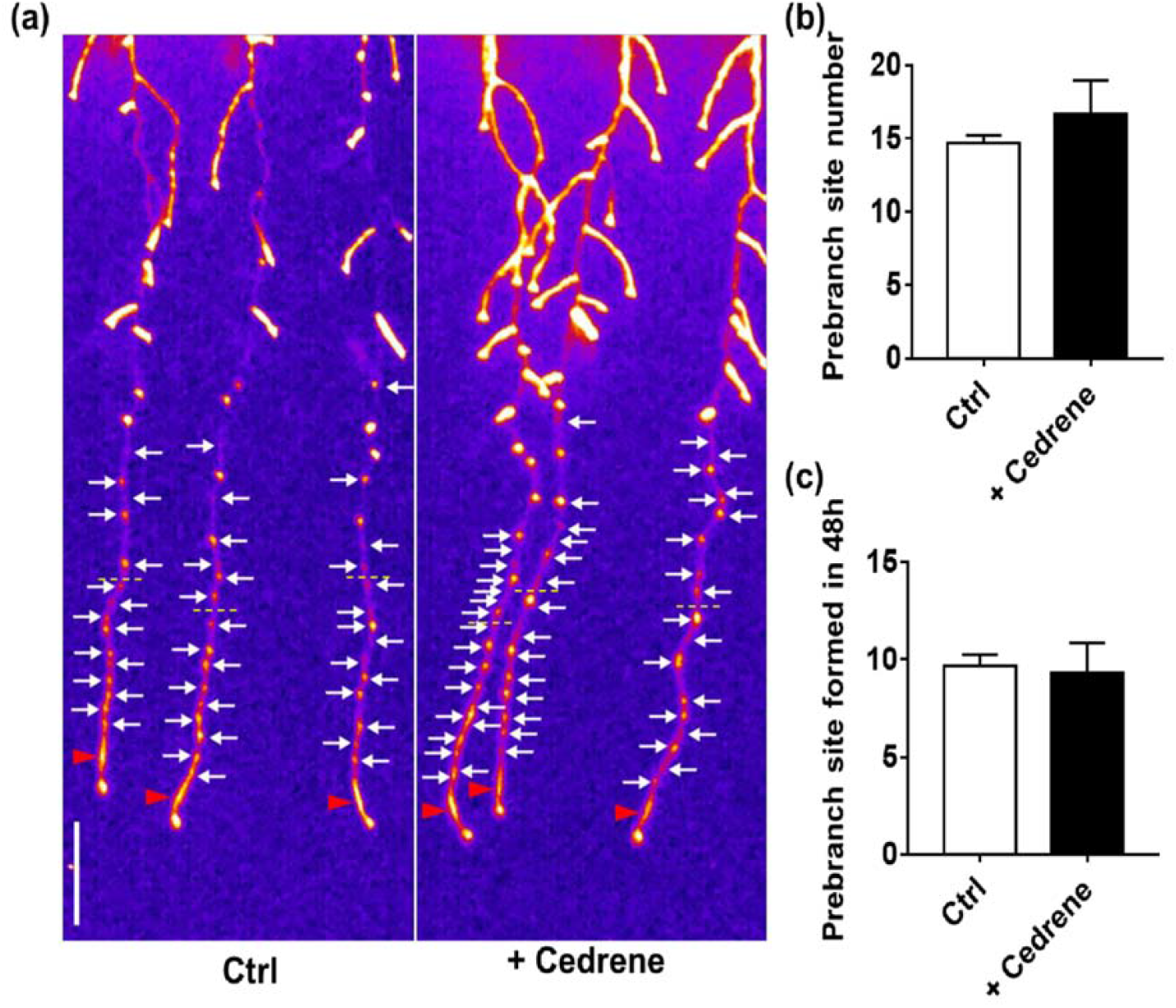
The effects of cedrene on prebranch site formation and *DR5* oscillation. (a and b) Luciferase imaging and quantification of prebranch site number of 3-day-old *pDR5:Luciferase* seedlings grown for 6d on medium supplied with or without 100 μM cedrene. The prebranch site in the newly formed primary root after transfer was measured. The red triangle indicates *pDR5:Luciferase* signal in the OZ, and white arrow indicates prebranch site revealed by persistent *pDR5:Luciferase* signal. Yellow dotted line indicates the position of root tip after 4 d co-cultivation. Scale bar, 1 cm. (c) Quantification of prebranch site number of *pDR5:Luciferase* seedlings within 48 h under control and cedrene-treated (100 μM) conditions. n ≥ 10. **, P < 0.01; *, P < 0.05 (Student’s *t*-test). Error bars indicate ± SD of the mean.

We did not observe differrence in the number of prebranch sites under cedrene treatment comparing with control treatment (Fig. 5b), which is in accordance with the observation of total LRP number under cedrene treatment (Fig. 4c). Since the periodicity and signal intensity of *DR5* oscillation are two important factors required for prebranch site formation, we further examined the effects of cedrene on those. We measured the periodic production of the prebranch sites in 48 h and found that cedrene treatment did not affect the formation of prebranch sites (Fig. 5c), indicating the periodicity of DR5 oscillation remain unchanged between control and cedrene treatments. Also, we did not observed a change in the expression intensity of the DR5 in OZ for plants grown in the presence of cedrene (Fig. 5a). Therefore, cedrene induces LR formation is not through affecting DR5 oscillation.

### Effect of cedrene on root development of auxin-related *Arabidopsis* mutants

Since cedrene can enhance the auxin response of early LRP, implying that auxin plays a role in cedrene-induced lateral roots formation, we further analyzed the response of wild-type (Col-0) and *Arabidopsis* mutants deficiented in genes related to auxin transport or response (*tir1afb2, slr-1/iaa14, arf7arf19, aux1-7, pin2, pin3*) to cedrene treatments (Fig. 6a). Col-0 and mutant lines were grown in medium supplemented with the solvent only or with 100 μM cedrene, and LR formation were analyzed after 8 d treatment. The results showed that in all these auxin signaling mutants, including *tir1afb2*, in which oscillation amplitude and prebranch site are drastically compromised (Xuan et al., 2015), *arf7arf19* and *slr-1/iaa14*, which completely abolished LR formation (Fukaki et al., 2002; Okushima et al., 2007), cedrene treatment did not influence LR formation and primary root elongation as compared with the control treatment (Fig. 6b, c). Moreover, the LR formation and primary root elongation promoting effect of cedrene on the auxin influx mutant *aux1-7* and auxin efflux mutant *pin2* were also disappeared (Fig. 6b, c). By contrast, the promoting effects of cedrene on LR formation and primary root elongation were not affected in the auxin efflux mutant *pin3*. These results suggested that cedrene-triggered LR formation and primary root elongation operates via a canonical auxin-response pathway and auxin transport system is necessary for that.

**Figure 6.**
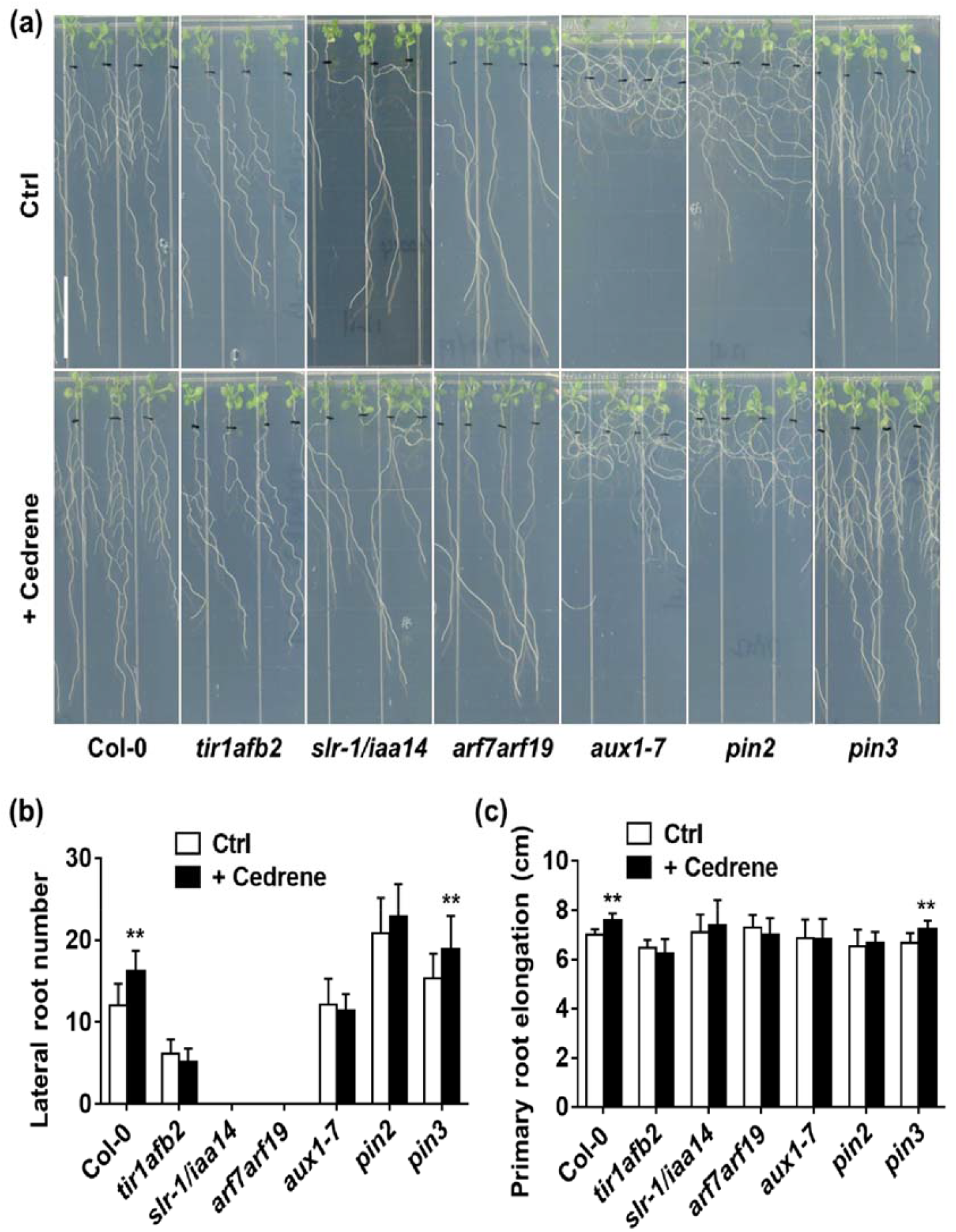
Influence of auxin signaling and transport on root system architecture modified by cedrene. (a) Representative photographs of wild-type (Col-0) and mutant seedlings of auxin signaling (*tir1afb2, slr-1/iaa14* and *arf7arf19*) and transport (*aux1-7, pin2* and *pin3*) under control and cedrene (100 μM) treatments for 8 d. Scale bar, 2 cm. (b and c) Quantification of emerged lateral root number (b) and primary root elongation (c) of plants shown in (a). n = 12. **, P < 0.01; *, P < 0.05 (Student’s *t*-test). Error bars indicate ± SD of the mean.

To further address the role of polar auxin transport in cedrene-mediated LR formation, LR formation was quantified in the presence of the polar auxin transport inhibitor 1-N-naphthylphthalamic acid (NPA) under control and cedrene-treated conditions (Fig. 7a). At 1 μM NPA, the ability of cedrene to stimulate LR formation was basically unaffected, whereas in the presence of 3 and 5 μM NPA, LR formation in response to cedrene was completely abolished (Fig. 7c). Moreover, the stimulation effect of cedrene on primary root elongation was completely abolished in the presence of NPA (Fig. 7b). Thus, it can be concluded that a functional auxin efflux machinery is required for cedrene-induced LR formation.

**Figure 7.**
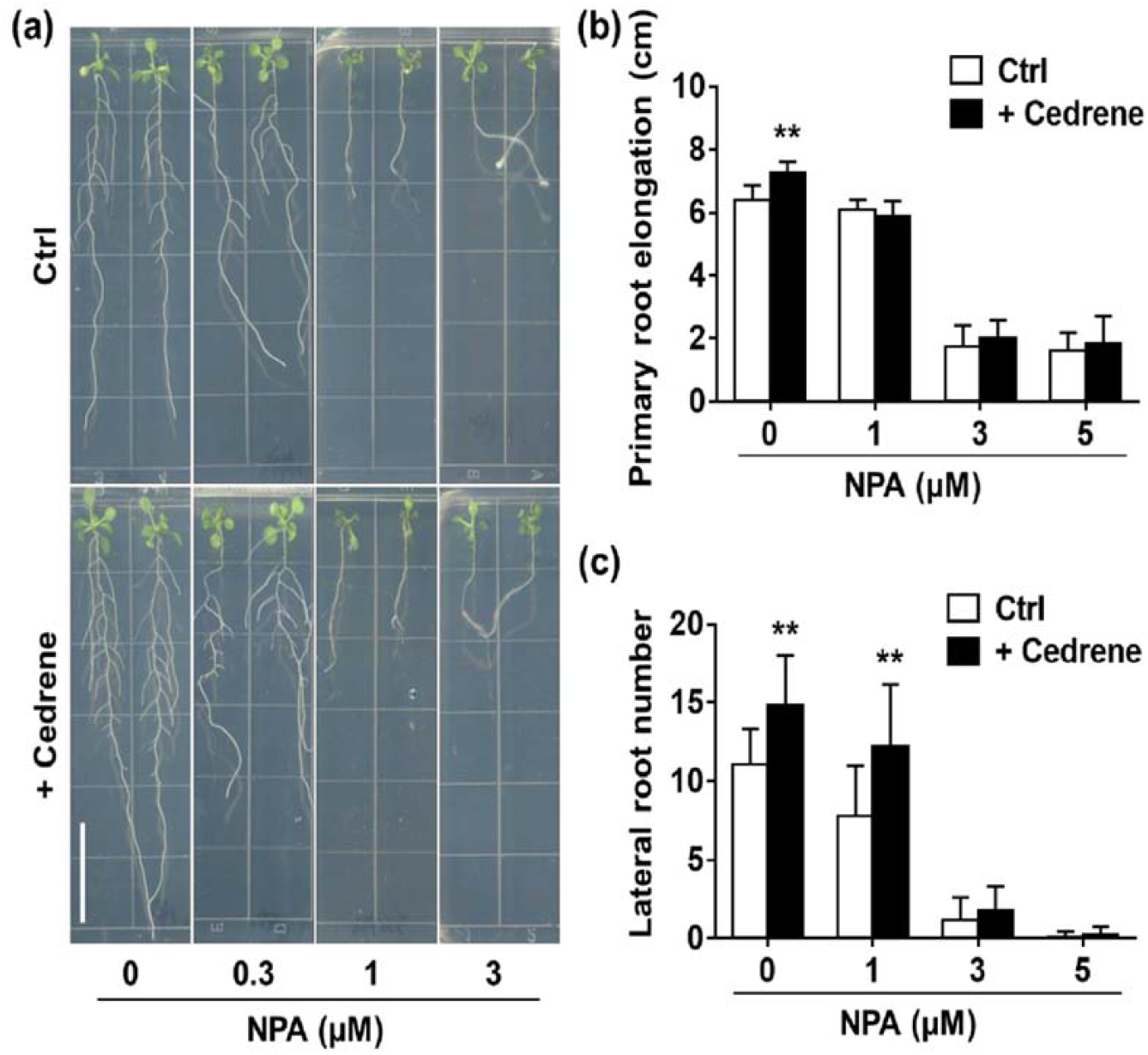
Effects of the polar auxin transport inhibitor NPA on cedrene-induced primary root elongation and LR formation. (a) Representative photographs of *Arabidopsis* seedlings grown under solvent (Ctrl) and cedrene (100 μM) treatments in presence of varying concentrations of NPA. Scale bar, 2 cm. (b and c) Quantification of emerged lateral root number (c) and primary root elongation (b) of plants shown in (a). n = 12. **, P < 0.01; *, P < 0.05 (Two-way ANOVA). Error bars indicate ± SD of the mean.

Since the polar auxin transport is mediated by polarly localized PIN proteins (van Berkel et al., 2013), we analyzed the expression pattern of PIN1, PIN2, PIN3 and PIN7 in primary root tips of seedlings expressing *pPIN1:PIN1:GFP, pPIN2:PIN2:GFP, pPIN3:PIN3: GFP* and *pPIN7:PIN7:GFP* (Fig. 8a-d), to test whether cedrene could regulate LR formation and/or primary root elongation through differential expression of the PINs family of auxin transporters. The results showed that the expression of *pPIN1:PIN1:GFP*, which was only detected in the stele of primary roots under control conditon, and *pPIN7:PIN7:GFP*, which was detected in the stele and columella of primary roots under control conditon, were not affected by cedrene treatment (Fig. 8a, e, d, h). By contrast, the expression of *pPIN2:PIN2:GFP*, which was detected in the cortex and epidermal cells under control conditon, was significantly increased when treated with cederen (Fig. 8b, f). Furthermore, the expression of *pPIN3:PIN3: GFP*, which was detected in the stele and columella of primary roots under control conditon, displayed a weak expression in the stele when treated with cedrene (Fig. 8c, g). These findings suggest that cedrene affects the expression and distribution of the PIN proteins in primary roots and the root response to cedrene does not occur in all tissues, but shows a clear preference for specific tissues and transport components.

**Figure 8.**
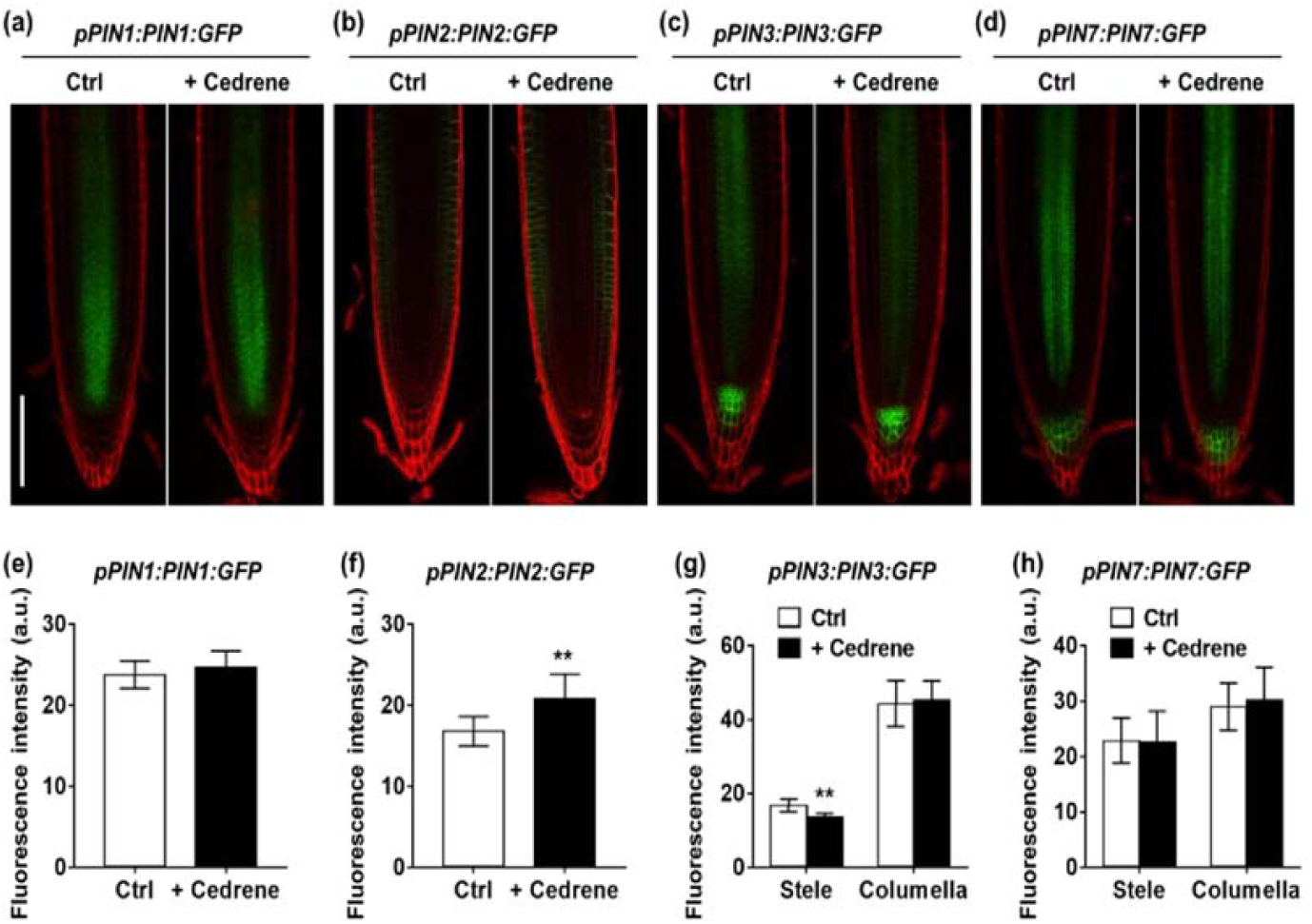
Expression of auxin efflux transporters in response to cedrene in primary roots. (a-d) Confocal images of *pPIN1:PIN1:GFP* (a), *pPIN2:PIN2:GFP* (b), *pPIN3:PIN3:GFP* (c) and *pPIN7:PIN7:GFP* (d) signal in the primary root tips of 3- day-old seedlings grown with or without cedrene for 6 d. The seedlings were stained with propidium iodide and analyzed by confocal microscopy. Scale bar, 100 μm. (e-h) Quantification of *pPIN1:PIN1:GFP* (e), *pPIN2:PIN2:GFP* (f), *pPIN3:PIN3:GFP* (g) and *pPIN7:PIN7:GFP* (h) signal intensity at primary root tip shown in (a-d). n ≥ 8. **, P < 0.01; *, P < 0.05 (Student’s *t*-test). Error bars indicate ± SD of the mean.

## DISSCUSSION

Previous studies on rhizosphere microbe’s beneficial function usually emphasized the first step of rhizospheric colonization, resulting in a direct contact with the root (Bais et al., 2004; Beauregard et al., 2013). However, volatile compounds (VCs) produced by microbes can spread to distant places through the air, porous soil, and liquid environment, so it should be an ideal signal substance to cross-exchange among different organisms(Maffei et al., 2011). Consistent with this, a comparative analysis of experimental data has shown that volatile metabolites made an enormous contribution to the microbial interactions than non-volatile ones (Tirranen and Gitelson, 2006). Our results clearly demonstrated that the exchange of molecules between *T. guizhouense* and *Arabidopsis* via intimate contact is not necessary (Fig.1a), and *T. guizhouense* VCs can act as signals to modulate plant growth and development.

Actually, it is well recognized that plant-associated microorganisms produce a large number of VCs, which have the potential to play an important role in mediating the interaction between plants and microbes (Kanchiswamy et al., 2015; Piechulla et al., 2017; Fincheira and Quiroz, 2018). So far, most of the effects mediated by microbial VCs have been obtained through co-culture experiments, that is, plants are exposed to complex mixtures of inorganic and organic volatiles released by microorganisms. This makes it difficult to disentangle the functions of individual compounds. On the other hand, the substrate availability and metabolic activities of the microorganisms also will affect the composition of microbial blends to a large extent (Fiddaman and Rossall, 1994). Regarding plants, the observation results obtained by different co-culture experiments vary greatly from strong growth inhibition to significant growth promotion (Splivallo et al., 2007; Li et al., 2021). In order to confirm and provide functional evidence for the microbial VCs action potential, it is necessary to identify distinct biologically active compounds and test them separately or in prescribed mixtures (Piechulla et al., 2017). Here, we discovered cedrene, a high-abundant SQT in the emission profile of *T. guizhouense* NJAU 4742, was sufficient to stimulate LR formation. Cedrene was previously identified in ascomycete *Muscodor albus*, and has the potential to kill a broad range of plant- and human-pathogenic fungi and bacteria synergistically with other sesquiterpenes volatiles (Strobel et al., 2001). Its function as a signaling compound in root branching as demonstrated in this study, has not been reported before. This evidence is closely related to the results previously reported by Ditengou et al. (2015) that (–)-thujopsene, as a low-abundant SQT, can also stimulate lateral root formation. Nevertheless, not all SQTs can induce lateral roots (Ditengou et al., 2015), indicating that the plant response to these chemicals is specific. Furthermore, along with other environmental factors, such as water, temperature and light, various distinct bioactive microbial volatiles that can affect root system architecture have been identified in bacteria and fungi (Bailly et al., 2014; Ditengou et al., 2015; Garnica-Vergara et al., 2016; Perez-Flores et al., 2017), emphasizing the high plasticity of plant roots in response to heterogeneous macro- and micro-conditions.

The phytohormone auxin fulfils multiple roles throughout LR development (Du and Scheres, 2018). Tryptophan treatment initiated additional LRPs were mostly originated from the auxin-producing pericycle sectors, suggesting that a local auxin input is able to specify lateral root founder cells (LRFCs) (Dubrovsky et al., 2008). In this study, an enhanced expression of *DR5*, which is an established marker for auxin response and indirectly for auxin accumulation (Ulmasov et al., 1997; Sabatini et al., 1999), was observed at early stages (I-III) LRPs and in the columella and meristematic protoxylem pole of root tip under cedrene treatment (Fig. 4a and b). These results suggest that cedrene promotes the formation of early stage LRP by regulating auxin homeostasis in LRPs. Before LRP initiation, the cell files of the xylem pole pericycle (XPP) at one side of the root will be specified and activated to form LRFCs, so that to regulate the spatial distribution of LRPs (Van Norman et al., 2013; Du and Scheres, 2018). The auxin-regulatory GATA23 transcription factor is considered as the first molecular marker for specification of LRFCs (De Rybel et al., 2010). Using a GATA23 RNAi line, in which the expression of GATA23 was reduced to about 30% of normal levels, a strong reduction in the number of stage I-II primordia and an overall decrease in the number of emerged primordia was observed. In contrast, overexpression of GATA23 resulted in a strong increase of stage I and II primordia (De Rybel et al., 2010). This result is basically consistent with our data, cedrene treatment enhanced the expression of GATA23 in root and increased the number of very early stage LRPs (stage I) and ELR, although the total number of LRP was not affected by cedrene treatment (Fig. 4b-d). These results imply that cedrene may stimulate the initiation of LRP by controlling founder cell identity of XPP cells via enhancing GATA23 expression. SOLITARY-ROOT (SLR)/IAA14-ARF7/ARF19 module is an important auxin signal component for LR initiation, in which auxin-induced degradation of unstable SLR protein that de-repressed ARF7 and ARF19 transcription factors, thus activating downstream gene expression (Fukaki et al., 2002; Fukaki et al., 2005; Okushima et al., 2005). Our results showed cedrene treatment could not rescue the lack of LRs in *slr-1/iaa14* and *arf7arf19* mutants and failed to induce LRs formation in the auxin perception double mutant *tir1afb2* (Fig. 6a, b). These results suggest that canonical auxin-response pathway is indispensable for cedrene induced lateral root formation.

In plants, auxin is generally transported by two types of carriers (i.e. influx carriers and efflux carriers). So far, some transmembrane proteins, such as AUX1/LIKE AUX1 (AUX1/LAX) family, with specific auxin influx functions have been described in *Arabidopsis* (Bennett et al., 1996; Marchant et al., 2002; Swarup et al., 2008). For auxin efflux, PIN protein family and ATP-binding cassette subfamily B (ABCB)-type transporters of the multidrug resistance/phosphoglycoprotein (ABCB/MDR/PGP) protein family have been identified to play the roles (Galweiler et al., 1998; Noh et al., 2001). Both auxin influx and efflux carriers can affect the formation of lateral roots (Marchant *et al*., 2002; Swarup *et al*., 2008; Peret *et al*., 2013). The LR reprogram ability of VCs produced by microbes does not only influence auxin signaling but also its transport through the root system. In *T. atroviride*, 6-PP enhanced the expression of the auxin transporters PIN1, 2 and 3 in the primary root of *Arabidopsis* (Garnica-Vergara et al., 2016). In *F. oxysporum*, the enhanced LR formation by VCs is abolished in auxin transport mutants (Bitas et al., 2015). Moreover, stimulation of LR in *Arabidopsis* by the ectomycorrhizal fungus *L. bicolor* requires PIN2-mediated auxin transport (Felten et al., 2009). Consistent with these observations, treatment with the polar auxin transport inhibitor NPA compromised the ability of cedrene to promote LR formation, suggesting that cedrene is subjected to polar transport. Furthermore, stimulation of LR formation by cedrene was completely abolished in mutant *aux1-7*, suggesting that cedrene-triggered LR formation also requires a complete auxin influx system.

The cedrene-induced LR formation and primary root elongation shown here opens new avenues for biotechnological application of microbial VCs, and demonstrates clearly that the operation of distinct bioactive microbial VCs is a practical tool to decode the underlying cellular and molecular reactions and mechanism occurring in microbial VC-mediated interactions. Understanding those mechanisms will be a prerequisite for the development of strategies for applying microbial VCs in plant growth in the future.

## MATERIALS AND METHODS

### Plant materials and growth conditions

*Arabidopsis thaliana* accessions Col-0 is the wild-type genotype. The marker lines *pDR5:GUS* (Ulmasov et al., 1997), *pDR5:Lucifease* (Moreno-Risueno et al., 2010), *pCYCB1;1:GUS* (Himanen et al., 2002), *pGATA23:nls-GUS* (De Rybel et al., 2010), *pPIN1:PIN1:GFP* (Benkova et al., 2003), *pPIN2:PIN2:GFP* (Blilou et al., 2005), *pPIN3:PIN3:GFP* (Zadnikova et al., 2010), *pPIN7:PIN7:GFP* (Blilou et al., 2005) and the mutant lines *tir1afb2* (Dharmasiri et al., 2005), *slr-1/iaa14* (Fukaki et al., 2002), *arf7arf19* (Okushima et al., 2007), *aux1-7* (Pickett et al., 1990), *pin2* (Roman et al., 1995), *pin3* (*salk_005544*) were used in this study. After 2-3 d of stratification at 4°C in the dark, *Arabidopsis* seeds were surface-sterilized with 30% (v/v) NaClO solution for 10 min. The seeds were germinated and grown on agar plates containing Murashige and Skoog Basal Salts Mixture (MS salts, PhytoTech LABS) in square Petri plates (10 ×10 cm). Standard growth medium consisted of 0.5 × MS salts (2.15 g l^-1^), 0.1g l^-1^ Myo-inositol, 0.5g l^-1^ 2-(N-morpholino) ethanesulfonic acid (MES), 1% sucrose (pH 5.7), and 1% Agar (Solarbio). Plants were vertically placed at an angle of 65° in a plant growth chamber, under a long-day photoperiod (16 h: 8 h, light: dark), with a light intensity of 100 μmol m^-2^ s^-1^, at 22 °C. After 3 d of growth, the seedlings were applied for further experiments.

### Growth of *T. guizhouense* NJAU 4742 and co-cultivation with plants

For experiments involving fungal VCs, bi-compartmented Petri dishes (9 cm diameter) were used. One of the compartments was filled with MS salts agar medium, and the other one was placed with a patch of MS salts agar medium (1.5 cm×1.5 cm) incubated with 3 ul *T. guizhouense* NJAU 4742 (maintained in the Jiangsu Provincial Key Lab for Organic Solid Waste Utilization, China) spores solution. After 3-4 d of fungal growth at 28°C, 3-day-old seedlings (4 plants) were transferred to the compartment filled with MS salts agar. The plates were sealed with breathable tape (MBT, a pressure-sensitive adhesive type usually used in medicine) and incubated for 8 d in a growth chamber at 22°C. At the end of this period, primary root length, lateral root number, and biomass production were recorded.

### Analysis of *T. guizhouense* NJAU 4742 VCs

*T. guizhouense* NJAU 4742 was grown in a head-space bottle containing MS salts agar medium for 6 d at 28 °C. At a constant temperature of 40°C, the head-space bottle was shaken for 60 min at a shaking speed of 450 rpm (5s on and 2s off). Then solid-phase microextraction (SPME) fiber (50/30 μm DVB/CAR on PDMS) was inserted into the headspace of the sample, and the sample was extracted in the headspace for 60 min. The SPME fiber was removed and desorbed at 250°C for 5min, and then was separated and identified by gas chromatography-mass spectrometry (GC-MS). GC-MS analysis was carried out by Agilent 7890B-5977B (GC-MS)_PAL RSI 120, equipped with a chromatographic column (Agilent DB-wax, 30m×0.25mm×0.25µm). Helium (> 99.999%) was used as the carrier gas (1.0 ml/min), and the injection temperature was 260 °C. The column was held at 40 °C for 3 min and then was programmed to increase by 4 °C per min to a final temperature of 220 °C, which was maintained for 10 min. The interface temperature, ion source temperature and quadrupole temperature was 280 °C, 230 °C and 150 °C respectively; the ionization mode was EI^+^, 70ev; the detector voltage was 901V; the scanning mode was full-scan, and the mass range was 20-650 (m/z). These compounds were identified by comparison with mass spectra from a library of NIST (https://webbook.nist.gov/chemistry/). The identification of cedrene was performed by comparing retention time (Rt) and the mass spectra from an authentic standard ((−)-α-cedrene, Sigma-Aldrich) with those obtained in the sample.

### Chemical preparation and treatments

1-N-naphthylphthalamic acid (NPA) and (−)-α-cedrene (≥ 95%, Sigma-Aldrich) were dissolved in DMSO to make a 50 mM and 200 mM stock solution, respectively. For treatment, the required amount of the stock solutions was added into MS salts agar and mixed in uniform before being poured into Petri dishes. For lovastatin treatment, lovastatin (PHR1285, Supelco), was dissolved in ethanol to make a 100 mM stock solution, sterile-filtered and added to the MS salts media at the indicated concentration. The plants and *T. guizhouense* were cultivated in a square bi-compartment Petri dish (10 cm × 10 cm), in which a small Petri dish (3cm diameter) was placed to inoculate *T. guizhouense. T. guizhouense*, not *Arabidopsis*, grown with lovastatin. LR formation and primary root length was measured after treatment for 7 days.

### Phenotypic data analysis

After co-cultivation with fungus or with cedrene for an indicated time period, the length of the newly elongated PR during the treatment was quantified and the emerged LRs in the whole PR were counted. The plates with seedlings were scanned using EPSON XL11000 for the measurement of PR elongation with Fiji software (http://fiji.sc/) and the emerged LRs were recorded under a microscope. The LR density was determined by dividing the emerged LR number by the new formed PR length for each analyzed seedling. The biomass production of each seedling was measured on an analytical balance. LRPs were quantified 6 d after co-cultivation. The *pCYCB1;1:GUS* seedlings were stained and cleared to visualize the LRPs at early stages of development and each LRP developmental stage was classified according to Malamy and Benfey (1997) as follows. Stage I, LRP initiation, in the longitudinal plane, approximately 8 to 10 “short” pericycle cells are formed. Stage II, the formed LRP is divided into two layers by a periclinal division. Stage III, the outer layer of the primordium is divided periclinally, generating a three-layer primordium. Stage IV, LRP with four cell layers. Stage V-VII, the midway between the LRP departing from the parent cortex to the LRP appears to be just about to emerge from the parent root.

### GUS histochemical staining

For histochemical analysis of GUS activity, the roots of 3-day-old *pDR5:GUS, pCYCB1;1:GUS* and *pGATA23:nls-GUS* marker lines were incubated overnight at 37 °C in a GUS reaction buffer after 6 d of cedrene treatment. The stained roots were cleared using the method of Malamy and Benfey (1997). For each marker line and each treatment, at least 8 transgenic plants were analyzed. A representative sample was chosen and photographed using Leica DM2500 microscope.

### Luciferase assay

*pDR5:Luciferase* expression along the primary root was analyzed by using Lumazon (Xuan et al., 2018). After 6 d of treatment with cedrene, *pDR5:Luciferase* plants were sprayed with a 1mM potassium luciferin (Gold Biotechnology) and reacted for 15 min in darkness, then were imaged immediately with a 15 min exposure time. For investigating the effects of cedrene on the periodicity of *DR5* oscillation, the position of root tip was marked after 4 d of co-cultivation and the prebranch sites formed in the following 2 d was counted. The picture series were saved as TIFF format by IVScopeEQ software for further analysis in Fiji (http://fiji.sc/). The Fiji lookup tables “Fire” was used to convert black and white images into color scales based on pixel intensity.

### Fluorescence microscopy

For confocal microscopy, control or cedrene-treated transgenic *Arabidopsis* seedlings (*pPIN1:PIN1:GFP, pPIN2:PIN2:GFP, pPIN3:PIN3:GFP* and *pPIN7:PIN7:GFP*) were mounted in 10 mg ml^-1^ propidium iodide solution on microscope slides. A Leica SP8 laser-scanning microscope was used for fluorescence imaging of the *Arabidopsis* roots. Each sample was analyzed separately for propidium iodide (with a 568-nm wavelength argon laser for excitation, and an emission window of 585–610 nm) and GFP fluorescence (488 nm excitation/505–550 nm emission). More than eight independent seedlings were analyzed per line, and treatment representative images were selected for figure construction.

